# Genomic sequence and structural variations accumulating between laboratory lineages of wild type *C. elegans*

**DOI:** 10.64898/2026.04.27.720957

**Authors:** Zachary D. Bush, Alice F. S. Naftaly, Devin Dinwiddie, Kenneth J. Hillers, Diana E. Libuda

## Abstract

Laboratory cultivation subjects model organisms to selective pressure and genetic drift that can result in the accumulation of many genomic and phenotypic differences over time. The nematode *Caenorhabditis elegans* has been used for research since the 1970s, and studies comparing the N2 Bristol and CB4856 Hawaiian isolates provided foundational knowledge about metazoan genome evolution. Most comparative genomics studies have used these isolates because their long-term geographical isolation promoted a high degree of genomic divergence within the species. Further, there is growing evidence of phenotypic differences between laboratory lineages of each wild type isolate after repeated independent lab cultivation of these strains. To examine the genomic divergence between different laboratory lineages the Bristol and Hawaiian backgrounds, we first generated *de novo* genome assemblies of two Bristol and two Hawaiian lineages from Illumina and PacBio sequencing reads. Following genome assembly, we quantified Single Nucleotide Polymorphisms (SNPs), short insertion/deletions (indels), and genomic structural variants (SVs). Between laboratory lineages of the Bristol isolate, we identified 25,432 SNPs, 5,202 indels, and 441 SVs. When aligning laboratory lineages of the Hawaiian isolate, we identified 4,518 SNPs, 1,188 indels, and 387 SVs. For both sets of comparisons, we find that SNPs and indels are broadly enriched in introns and depleted from coding sequences. In contrast to SNPs and indels, we find that genomic SVs are enriched in intergenic sequences. Taken together, our analyses reveal the accumulation of genomic divergence between lineages of Bristol and Hawaiian *C. elegans* from independent lab cultivation, and how these variants may underpin emergent phenotypic differences observed in the two most popularly used *C. elegans* wild type isolates.

**Author Summary:** Laboratory model organisms, like natural populations, are subject to evolutionary pressures and genomic changes during prolonged laboratory cultivation. In this study we comprehensively quantify SNPs, indels, and SVs between independent lab cultivations of the *C. elegans* lineages of the Bristol and Hawaiian isolates.

## Introduction

DNA mutations are a source of genetic variation and the prerequisite for many evolutionary processes acting on populations. In the human genome, the rate and prevalence of single nucleotide polymorphisms (SNPs) are the highest contributors towards mutations, followed by small insertions/deletions (indels, < 50 bp) and structural variants (SVs, ≥ 50 bp), respectively (The 1000 Genomes Project Consortium 2012; Nesta, Tafur, and Beck 2021). Mutations, whether they affect one, thousands, or millions of base pairs, can have profound impacts on the development, health, and survival of individuals (Stankiewicz and Lupski 2010; Gagliano et al. 2019). Combining modern DNA sequencing technologies with the use of model organisms has greatly enhanced our detection and understanding of genomic variation in the contexts of both developmental and evolutionary biology.

Model organisms used in labs are also subject to the same processes of mutation and evolution. Genetic drift and repeated selection for desirable phenotypes in model organisms can result in lab-to-lab variation in the genetic background of wild type strains. Several studies in bacteria, yeast, plants, and vertebrates demonstrate that laboratory cultivation of model organisms leads to the accumulation of genomic variation with functional consequences on processes like metabolism and reproduction (Yalcin et al. 2004; Guryev et al. 2006; Alonso-Blanco et al. 2003; Bentsink et al. 2006; Daranlapujade et al. 2003; Bradley et al. 2016). Further, some inbred lines of laboratory mice have significant differences in their mutation rate (Chebib et al. 2021; Uchimura et al. 2015). Thus, understanding how genetic variation between different lineages of laboratory wild types may affect the practice and interpretation of some research results.

While much prior work have focused on characterizing genetic variation between wild type strains via the accumulation of SNPs and indels (Daranlapujade et al. 2003; Carreto et al. 2008; Y. Wang et al. 2018; Guryev et al. 2006), the extent and impact of large genomic SVs between the genomes of laboratory lineages of wild type models has remained largely elusive due to difficulties of earlier sequencing technologies in detecting and mapping these large, complex genomic regions. Current long-read sequencing technologies have greatly aided the detection of SVs in humans and revealed that SVs between human populations affect many more base pairs than SNPs and indels (Hurles, Dermitzakis, and Tyler-Smith 2008; Sudmant et al. 2015; Sedlazeck et al. 2018). Additionally Large, newer tools utilizing reciprocal whole-genome alignments of long-read *C. elegans* genome assemblies have both aided the detection of SVs that far exceed the size of sequencing reads and more complex SVs such as inversions, translocations, and duplications (Bush, Zachary D. et al. 2025; Goel et al. 2019). genomic SVs could pose disruptions to daily research practices if, for example, thousands of bases have been deleted or rearranged in an area targeted by PCR or CRISPR. Further, SVs are known to disrupt gene expression and lead to disease phenotypes such as cancer (Hurles, Dermitzakis, and Tyler-Smith 2008; Stankiewicz and Lupski 2010; Sakamoto et al. 2021). Knowing that disruptive SVs are likely accumulating in laboratory model organisms requires a more comprehensive analysis of genomic variation between laboratory model systems.

*Caenorhabditis elegans* is an excellent model organism to study the accumulation of mutations in laboratory model organisms. Many *C. elegans* researchers use the N2 Bristol isolate as the canonical wild type since it was first isolate established and extensively characterized in a laboratory setting by Sydney Brenner in the 1970s (Brenner 1974; Sulston and Brenner 1974). Many labs also use the CB4856 Hawaiian isolate to perform genetic mapping as well as to study genomic variation, recombination, and evolution. (Hodgkin and Doniach 1997; Koch et al. 2000; Wicks et al. 2001; Rockman and Kruglyak 2009; Thompson et al. 2015; Crombie et al. 2019). While the ∼30,000-50,000 generations of sequence divergence between the N2 Bristol and CB4856 Hawaiian wild types has been determined by multiple groups over the course of increasing quality and depth of sequencing technologies (Wicks et al. 2001; Swan et al. 2002; Seidel, Rockman, and Kruglyak 2008; Thompson et al. 2015; Kim et al. 2019; D. Lee et al. 2021; Bush, Zachary D. et al. 2025), recent work on germline mutation rates, suggest that considerable genetic variation has also accumulated within the N2 Bristol strain since laboratory domestication (Denver et al. 2009; Saxena et al. 2019). For *C. elegans,* the germline rate of mutation accumulation is approximately 2.7 to 3.0 × 10^−9^ mutations per site per generation (Denver et al. 2009; Saxena et al. 2019) and the generation time is approximately three days. Depending on how often labs return to frozen stocks, each laboratory N2 lineage alone may have accumulated up to ∼1,500 single nucleotide mutations since the 1970s and nearly 790 potential mutations since the first genome was published in 1998 (C. elegans Sequencing Consortium 1998). Notably, this predicted variation does not include the accumulation of multi-nucleotide indels and genomic SVs. While isolated examples of genomic variation in N2 between labs has been documented, a modern genome wide analysis has yet to reveal the full extent of genetic drift between labs.

The genomes of N2 Bristol and CB4856 Hawaiian that are currently in individual labs likely carry considerable genomic variation relative to other labs isolates. Previous studies using earlier genome assemblies have identified many segmental duplications between lab lineages of *C. elegans* wild type strains (Vergara et al. 2009) as well as duplications ranging in size from 200 bp to 108 kb that affect as many as 26 genes (Vergara et al. 2009). This variation may underpin phenotypic variation as well as previous work that has shown the lifespans of laboratory N2 Bristol isolates varies between 12-17 days (Gems and Riddle 2000). Further, prior research suggest that genetic variation acquired from laboratory use in wild type backgrounds contributes to many phenotypic differences in reproduction (10% fewer sperm and self-progeny), social versus solitary feeding, sensory signaling (CO_2_ avoidance), and social behaviors (>3-fold difference in rates of aggregation/bordering) (Sterken et al. 2015; Andersen et al. 2014; Duveau and Félix 2012; Weber et al. 2010; McGrath et al. 2009; Rogers et al. 2003). Given the growing evidence for phenotypic divergence between canonical wild type *C. elegans,* it is increasingly important to understand the underlying genomic changes that lead to these phenotypic differences.

Recently, several labs produced *de novo* genome assemblies for their lineages of the Bristol and Hawaiian isolates using a combination of Illumina, PacBio, and Oxford Nanopore sequencing platforms (Yoshimura et al. 2019; Kim et al. 2019; Ichikawa et al. 2024; Bush, Zachary D. et al. 2025). Compared to the previous short-read based assemblies of N2 Bristol (C. elegans Sequencing Consortium 1998; Hillier et al. 2005), the newer assembly of N2 Bristol from 2019, called VC2010 (Flibotte et al. 2010; Tyson et al. 2018; Yoshimura et al. 2019), identified 53 more predicted genes, 1.8 Mb of additional sequence, and eliminated 98% of existing gaps in the prior N2 Bristol reference genome. Thus, the overall structure of the VC2010 Bristol genome very likely represents the genome of Bristol *C. elegans* currently used in laboratories worldwide (Yoshimura et al. 2019). In 2019, the first *de novo* CB4856 Hawaiian assembly from long-read sequencing extended the length of the Hawaiian genome, which included over 3,000 previously uncharacterized SVs (Kim et al. 2019). Further, *de novo* genome assembly and comparison of both the N2 Bristol genome and CB4856 Hawaiian genome from highly accurate PacBio Hi-Fi reads comprehensively identified SVs and transposable element mobility between the two diverged isolates (Bush, Zachary D. et al. 2025). At present, the advent of multiple long-read genome assemblies of the Bristol and Hawaiian isolates from different labs provides the first opportunity for a modern analysis of SNPs, indels, and SVs that accumulate from laboratory cultivation of *C. elegans*.

To determine the extent of genetic variation between our laboratory lineages of N2 Bristol and CB4856 Hawaiian, we utilized short- and long-read alignments as well as reciprocal whole-genome alignments our DLW Bristol and DLW Hawaiian genome assemblies from our lab to the VC2010 Bristol (Yoshimura et al. 2019) and Kim CB4856 Hawaiian (Kim et al. 2019) genomes, respectively. We identified SNPs, indels, and SVs unique to the wild type strains in each lab. Between the different lab lineages of the Bristol strain, we identify over 724.9 kb of genomic variation. When aligning lab lineages of the CB4856 Hawaiian isolate, we identify over 2.07 Mb of sequence variation. Notably, 78.6% and 95.2% of the genomic variation in Bristol and Hawaiian genomes, respectively, are due to SVs. We find that SNPs and indels are largely enriched in introns, while SVs are often observed in intergenic sequences of the *C. elegans* genome. We do find some evidence for an enrichment of indels in several transcription factor binding sequences in the Bristol background, suggesting that variations in regulatory sequences and other non-coding regions may underlie the phenotypic variances previously observed between laboratory strains. Taken together, our analysis of genetic variation between wild type laboratory strains highlights the impact of small sequence variants, large structural variants, and other chromosomal rearrangements accumulating in the genomes of laboratory model organisms.

## Methods

### *C. elegans* culture and sucrose floatation

The DLW N2 Bristol and DLW CB4856 Hawaiian strains of *C. elegans* were grown at 20°C on standard NGM agar plates seeded with the OP50 strain of *E. coli* as a food source. The DLW N2 Bristol originated from cultivation of the N2 Bristol strain from the lab of Anne Villeneuve and brought to the lab of Diana Libuda in January 2015, frozen into the Libuda Lab collection on March 16, 2015(Bush, Zachary D. et al. 2025). VC2010 is the Moerman Gene Knockout Lab subculture of N2 (Flibotte et al. 2010; Yoshimura et al. 2019; Tyson et al. 2018). DLW CB4856 Hawaiian is cultivation of CB4856 Hawaiian obtained from the CGC on February 5, 2015 and frozen into the Libuda Lab collection on March 4, 2015 and thawed for immediate sequencing (Bush, Zachary D. et al. 2025). To minimize bacterial contamination in downstream gDNA sample preps, we performed sucrose floatation on pooled populations of each isolate. Worms were washed from plates with 8mL cold M9 buffer and transferred to 15 mL glass centrifuge tubes using a glass Pasteur pipette. Collected worms were centrifuged at 3000 rpm at 4°C and washed in 4 mL of fresh M9 twice. To separate worms from bacteria and other debris, 4mL of 60% sucrose solution was added to 4 mL of M9 buffer and worms and vortexed briefly. The mixture was then spun at 5000 rpm at 4°C for 5 minutes. Using a glass pipette, the floating layer of worms were transferred to a new glass centrifuge tube on ice and brought up to 4 mL in fresh M9. Worms were then incubated at room temp for 30 minutes and gently vortexed every five minutes. Worms were washed three times in equal volume of fresh M9 were performed before storing collected worms in M9 at 20°C before genomic DNA (gDNA) extraction.

### Bristol and Hawaiian genome assemblies

To ensure the most equal genomic comparisons, all genomes were assembled from similar sequencing platforms (PacBio and Illumina) and subjected to the same processes of assembly, scaffolding, and base correction. Prior to assembly, PacBio and Illumina reads from all genomes were filtered by removing any reads that aligned to any of the 3000 bacterial genomes downloaded from the European Nucleotide Archive (ftp://ftp.ebi.ac.uk/pub/databases/fastafiles/embl_genomes/genomes/Bacteria). The DLW Bristol and DLW Hawaiian genomes used were assembled *de novo* from PacBio HiFi CCS long reads. The VC2010 Bristol genome (European Nucleotide Archive: PRJEB28388) and the Kim CB4856 Hawaiian genome (NCBI BioProject PRJNA523481) were re-assembled *de novo* from PacBio CLR reads. Three contig sets were generated for each genome using Canu (Koren et al. 2017) (version 2.2). Three runs of Canu produced identical contig sets for the DLW Bristol and DLW Hawaiian genomes, so only one set from each genome was retained for the following assembly steps. Each contig set was then polished 5 times with their respective raw PacBio reads (CLR for VC2010 Bristol and Kim Hawaiian contigs, CCS reads for the DLW genomes) using Racon (Vaser et al. 2017). Further, each PacBio-corrected contig set was then subjected to five more rounds of polishing with each genome’s respective set of Illumina short reads using Pilon (Walker et al. 2014). Genomes were then scaffolded to the short-read N2 Bristol reference genome (cel235) using Minimap2 alignments (Li 2018) via RagTag (Alonge et al. 2019; 2022) to generate chromosome level assemblies. For the VC2010 Bristol and Kim CB4856 Hawaiian assemblies, we created a final consensus scaffold using RagTag Patch to fill non-overlapping gaps between each of scaffold assemblies made from the three independent contig sets. Finally, each chromosome level assembly was subjected to two final rounds of base corrections with each genome’s PacBio and Illumina reads using Racon and Pilon, respectively. For all future variant calling methods, the DLW Bristol and DLW Hawaiian genomes were used as the reference sequence. Genome assembly quality was evaluated using BUSCO (Manni et al. 2021; Simão et al. 2015) and QUAST-LG (Gurevich et al. 2013; Mikheenko et al. 2018). Data from these quality checks are provided in Supplemental Figure S1 and Supplemental Data Files S5-8, respectively.

### SNP and indel variant calling in Bristol and Hawaiian genomes

To call short sequence variants, Illumina short reads from the VC2010 Bristol and Kim Hawaiian genomes were aligned to the DLW Bristol genome and DLW CB4856 Hawaiian genome genomes, respectively. Reads were then aligned to DLW reference genomes using BWA-MEM (Li and Durbin 2009). Aligned reads in SAM format where coordinate sorted using SAMtools (Li et al. 2009) and converted to BAM files. Picard was then used to assign read groups via AddOrReplaceReadGroups, and duplicate reads were filtered using MarkDuplicates (“Picard Toolkit” 2019). BAM files with filtered reads and alignments with a minimum mapping quality score of 40 were used then to call SNP and indel variants using HaplotypeCaller from the Genome Analysis Toolkit (GATK) (McKenna et al. 2010; Poplin et al. 2018). SNPs and indels were then hard-filtered according to GATK’s guidelines using the SelectVariants tool with the filter expression ‘‘QD < 2.0 || FS > 60.0 || MQ < 40.0 || MQRankSum < −12.5 || ReadPosRankSum < −8.0 || SOR > 4.0.” VCF files of SNPs and indels for the Bristol and Hawaiian genome analyses are provided in Supplemental Data Files S1 and S2.

### Whole-genome alignment and calling genomic structural variants

All assembly-to-assembly alignments were performed using Minimap2 (Li 2018) and SyRI (Goel et al. 2019) was then used to parse SAM files to call SVs. DLW Bristol and DLW Hawaiian genomes were used as the reference sequence. SVs were retained if they were at least 50 base pairs in length. Synteny calculations were performed using SyRI outputs that detail blocks of synteny and alignments within each parent block. Unaligned bases in each comparison were calculated by summing the lengths of “NOTAL” blocks output for each DLW reference genome VCF files of SV calls are provided in Supplementary Data Files S3 and S4. Example visualizations of SV calls using the Integrative Genomics Viewer (IGV) (Robinson et al. 2011) are provided in Supplemental Figure S6.

### Converting gene annotations between assemblies

The coordinates of gene annotations from the original N2 reference assembly (cel235) to corresponding regions in each of the Bristol and Hawaiian. The gene annotations for the WBcel235 assembly were downloaded in GFF3 format from Ensembl (http://ftp.ensembl.org/pub/release-105/gff3/caenorhabditis_elegans/). Annotations for promoters, enhancers, and transcription factor binding sites in the cel235 assembly were downloaded from WormBase via JBrowse. Assembly-to-assembly alignments via Minimap2 and Liftoff (Li 2018; Shumate and Salzberg 2021) were used to remap annotations onto each of the four new genome assemblies in this study. Intergenic and intronic regions were calculated separately using the BedTools suite (Quinlan and Hall 2010) and included in final annotation files.

### Assessing enrichment or depletion of variants in gene annotations

SNPs, indels, and SVs were tested for their degree of association in gene annotations to determine if lab lineages of each strain were accumulating genetic variants in coding versus noncoding sequences. Log2(fold) values were computed using the Genomic Association Tester (GAT) tool (Heger et al. 2013). Briefly, the observed overlap of each variant type with annotations was calculated independently and then compared to the mean overlap of simulated null distributions of each variant type. Simulated distributions were made from 20,000 iterations of a random uniform distribution across each chromosome. Input BED files for SNP, indel, and SV intervals were made by exporting pandas data frames of each VCF in python as a tab-separated file. GFF3 formatted files were then similarly converted to BED format. Statistical significance of each fold difference was determined via hypergeometric tests.

## Results

### *De novo* genome assembly from four laboratory lineages of wild type *C. elegans*

To enable the most accurate comparisons between each genome, we used the same pipeline for the genome assembly of each laboratory lineage of either the Bristol (VC2010 Bristol and DLW N2 Bristol; Yoshimura et al. 2019; Bush, Zachary D. et al. 2025) or Hawaiian isolates (Kim CB4856 Hawaiian and DLW CB4856 Hawaiian; Kim et al. 2019; Bush, Zachary D. et al. 2025). For the Bristol and Hawaiian genome sequencing and assembly performed by our lab, we denote these lineages as “DLW Bristol” and “DLW Hawaiian”. PacBio reads (HiFi CCS technology for DLW Bristol and Hawaiian (Bush, Zachary D. et al. 2025), CLR technology for VC2010 Bristol (Tyson et al. 2018; Yoshimura et al. 2019) and Kim Hawaiian (Kim et al. 2019) were used to generate contigs via the Canu assembler (Koren et al. 2017) after removal of all sequencing reads that align to any bacterial genomes (0.02% reads for DLW N2 Bristol; 0.1% reads for DLW CB4856 Hawaiian; 2.6% reads for VC2010 N2 Bristol; 6.3% reads for Kim CB4856; see Methods). Contigs for each genome were then subjected to multiple rounds of correction using alignments of both long and short reads (see Methods). All contigs were scaffolded into chromosome-level assemblies based on homology to a previous N2 Bristol assembly (C. elegans Sequencing Consortium 1998) followed by two final rounds of base corrections via read alignments. The DLW Bristol genome was assembled from 59 contigs to a final length of 105.2 Mb, with an average sequencing coverage of 513X for Illumina reads and 133X for PacBio reads (Table 1, Figures 1A and 1E). In comparison, the VC2010 Bristol genome was assembled from 183 contigs to a final length of 103.1 Mb, with an average coverage of 92X for Illumina reads and 187X for PacBio reads (Table 1, Figures 1B and 1E). The 105.8 Mb DLW Hawaiian genome was assembled from 109 contigs with average coverage of 590X from Illumina reads and 127X from PacBio reads (Table 1, Figures 1C and 1E). Finally, the Kim Hawaiian genome was assembled in our pipeline to a final length of 100.9 Mb from 344 contigs using 72X coverage from Illumina reads and 155X coverage from PacBio reads (Table 1, Figures 1D and 1E). The individual differences in chromosome lengths between each genome assembly was roughly proportional to the differences in total genome length (Figure 1F), which suggested that differences in final genome sizes were not due to grossly misassembled chromosomes.

**Figure 1.**
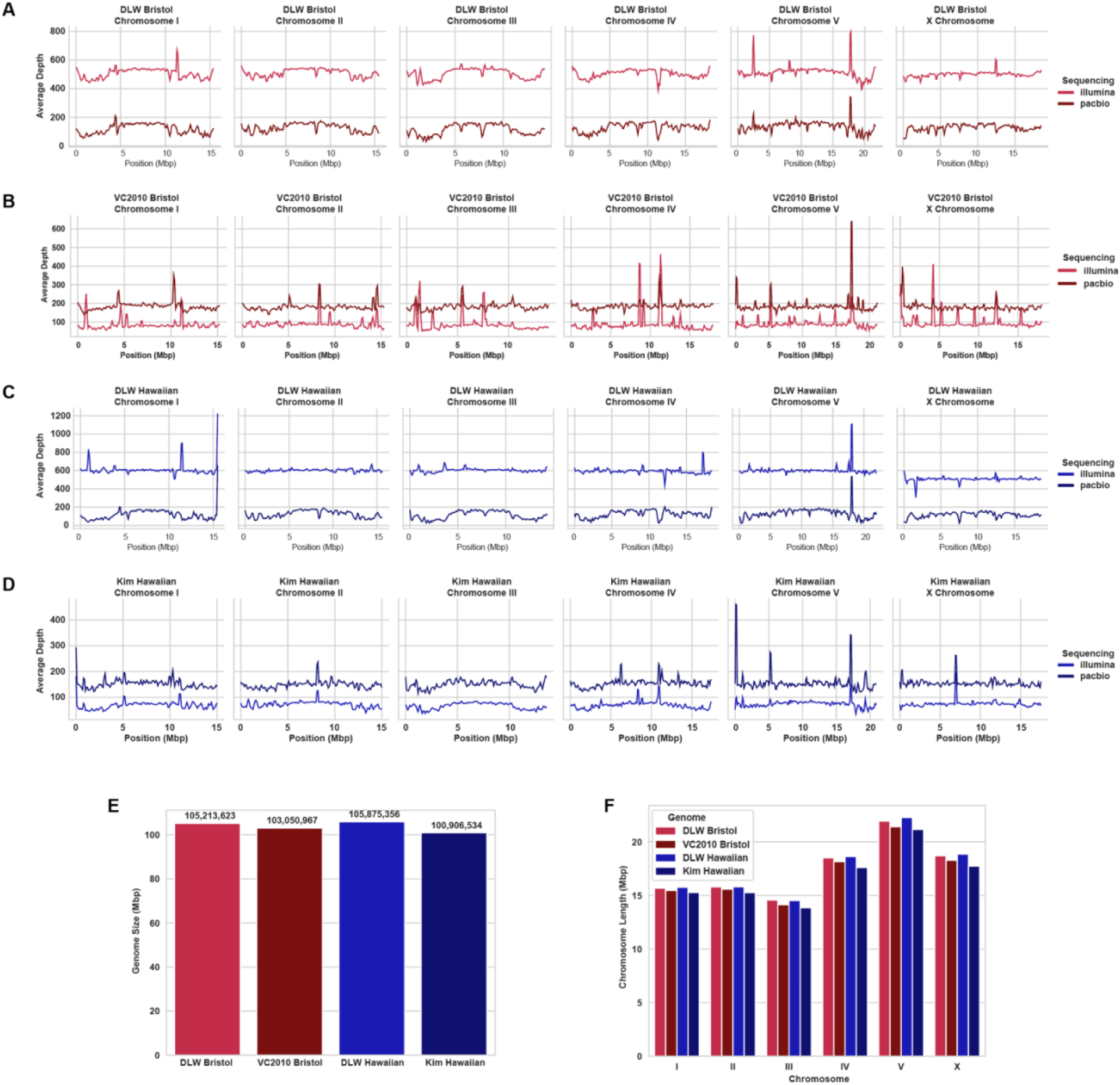
Genome assembly of four laboratory lineages of *C. elegans*. **(A)** Line plots showing the depth of sequencing coverage for PacBio and Illumina reads across each chromosome in the DLW Bristol genome assembly. **(B)** Line plots showing the depth of sequencing coverage for PacBio and Illumina reads across each chromosome in the VC2010 Bristol genome assembly. **(C)** Line plots showing the depth of sequencing coverage for PacBio and Illumina reads across each chromosome in the DLW Hawaiian genome assembly. **(D)** Line plots showing the depth of sequencing coverage for PacBio and Illumina reads across each chromosome in the Kim Hawaiian genome assembly. For all coverage plots, the average depth of coverage was calculated in a sliding 200 kb window with a 100 kb step size. **(E)** Bar chart depicting the total length of each assembled genome. **(F)** Bar chart showing the individual chromosome lengths in each Bristol or Hawaiian genome assembly.

**Table 1.**
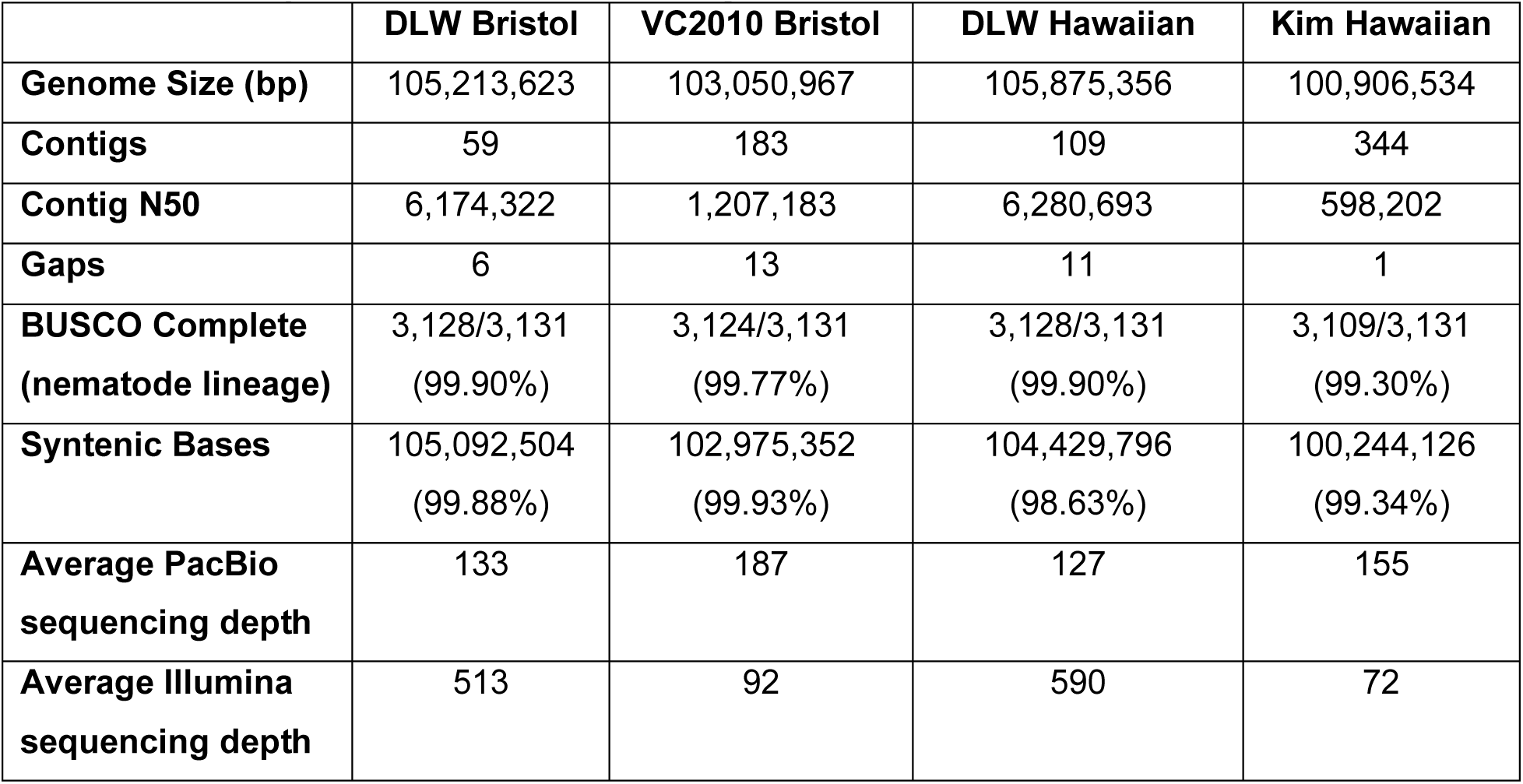
Summary of Bristol and Hawaiian genome assemblies.

To further check the quality of our genome assembly, we analyzed the presence of conserved gene orthologs and measured the degree of synteny between the pairs of Bristol and Hawaiian genomes. BUSCO analyses (Simão et al. 2015; Manni et al. 2021) reveals a high degree of complete gene orthologs (≥98.6% for all genomes) in the nematode lineage for all genome assemblies (Table 1, Supplemental Figure S1). Since each Bristol or Hawaiian lineage is derived from the same ancestral population, we expected a high degree of synteny, the co-linear organization of sequence content, in our genome comparisons. Indeed, for the Bristol lineages, we found that >99.8% of bases were syntenic with only 84.5 kb unable to be aligned in the genome. For the Hawaiian lineages, >98.6% of bases were found to be syntenic, but, in contrast to the Bristol lineages, there were 1.18 Mb that were unable to be aligned using our whole-genome alignment methods (see Methods; Goel et al. 2019). Given the high read coverage across all platforms in each genome (Table 1) and similar the Bristol lineages not sharing this degree of unalignable bases, this result may reflect the presence of genomic SVs as opposed to genome assembly errors. Taken together, these results indicate that our assembly of these four genomes are of sufficiently high quality and enable the accurate genomic comparisons between laboratory lineages of Bristol and Hawaiian *C. elegans*.

### Accumulation of short sequence variants in laboratory *C. elegans*

The Bristol and Hawaiian strains are estimated to have diverged from each other 30,000-50,000 generations ago (Thomas et al. 2015), and many studies have since documented the rich amount of genomic differences between these long-separated isolates (Maydan et al. 2010; Wicks et al. 2001; Thompson et al. 2015; Kim et al. 2019; Bush, Zachary D. et al. 2025). In contrast, since the ancestral stock of N2 Bristol was frozen in Sydney Brenner’s lab in 1968 (McCulloch and Gems 2001) and the discovery of the CB4856 Hawaiian isolate in 1972 (and a generation time of 3-4 days), a maximum of ∼5,000-7,000 generations could have passed for each background. Thus, we wanted to detect how much genomic sequence variation has accumulated within the Bristol or Hawaiian backgrounds during independent laboratory cultivation. Between the DLW Bristol and the VC2010 Bristol lineages, our analyses find that there are 25,432 SNPs and 5,202 indels, of which 1.07% of SNPs are homozygous and 3.42% of indels are homozygous (Table 2). Between the DLW Hawaiian and the Kim Hawaiian lineages, our analyses find that there are 4,518 SNPs and 1,188 indels, of which 1.79% of SNPs are homozygous and 4.12% of indels are homozygous (Table 2, Figure 2A-D). Thus, there may be roughly five times as many SNPs and indels between the two Bristol lab lineage genomes relative to our alignments of the two Hawaiian lab lineage genomes. In summary, many short sequence variants are accumulating through laboratory cultivation.

**Figure 2.**
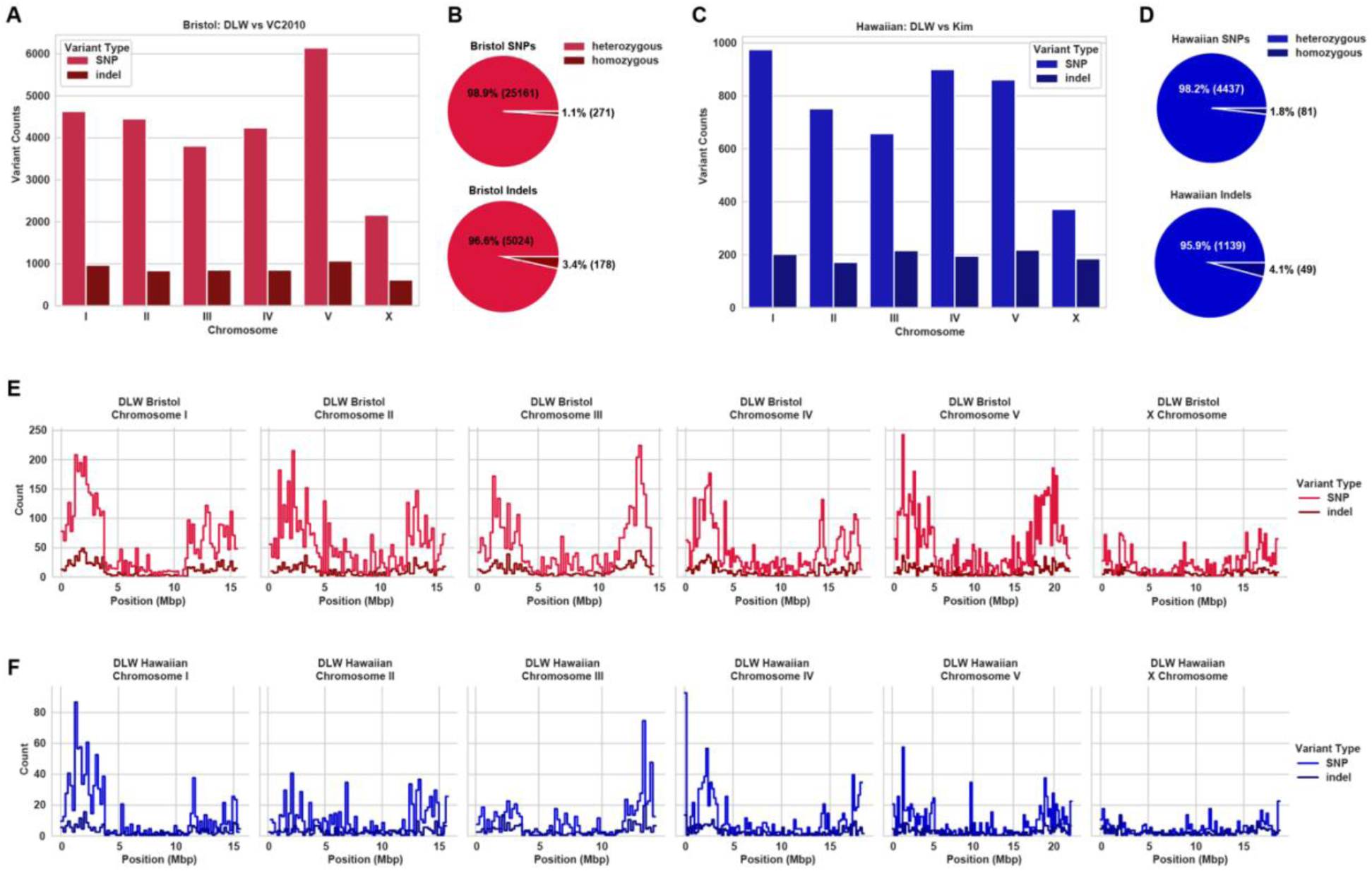
The genomic distribution of short sequence variants in Bristol and Hawaiian lineages. **(A)** Bar chart showing the number of SNPs and indels between the DLW Bristol and VC2010 Bristol genomes. **(B)** Pie charts showing the fractions of SNPs and indels in the Bristol genome comparisons that are heterozygous versus homozygous variant sites. **(C)** Bar chart showing the number of SNPs and indels between the DLW Hawaiian and Kim Hawaiian genomes. **(D)** Pie charts showing the fractions of SNPs and indels in the Hawaiian genome comparisons that are heterozygous versus homozygous variant sites. **(E)** Histograms showing the counts of SNPs and indels in non-overlapping 200 kb bins in the Bristol genomes. **(F)** Histograms showing the counts of SNPs and indels in non-overlapping 200 kb bins in the Hawaiian genomes.

**Table 2.** Summary of genomic variation between lineages of Bristol and Hawaiian *C. elegans*.

|  | Chromosome |  |  |  |  |  | Total |
| --- | --- | --- | --- | --- | --- | --- | --- |
|  | I | II | III | IV | V | X |  |
| **Bristol SNPs**<br>**(homozygous + heterozygous)** | 4,631 | 4,453 | 3,805 | 4,239 | 6,143 | 2,161 | 25,432 |
| **Bristol SNPs**<br>**(homozygous)** | 41<br>(0.89%) | 33<br>(0.74%) | 35<br>(0.92%) | 58<br>(1.37%) | 60<br>(0.98%) | 44<br>(2.04%) | 271<br>(1.07%) |
| **Bristol Indels**<br>**(homozygous + heterozygous)** | 970 | 839 | 853 | 853 | 1,070 | 617 | 5,202 |
| **Bristol Indels**<br>**(homozygous)** | 26<br>(2.68%) | 26<br>(3.10%) | 27<br>(3.17%) | 40<br>(4.69%) | 25<br>(2.34%) | 34<br>(5.51%) | 178<br>(3.42%) |
| **Bristol SVs** | 78 | 62 | 77 | 71 | 77 | 76 | 441 |
| **Hawaiian SNPs**<br>**(homozygous + heterozygous)** | 975 | 752 | 658 | 900 | 861 | 372 | 4,518 |
| **Hawaiian SNPs**<br>**(homozygous)** | 5<br>(0.51%) | 4<br>(0.53%) | 5<br>(0.76%) | 53<br>(5.89%) | 2<br>(0.23%) | 12<br>(3.23%) | 81<br>(1.79%) |
| **Hawaiian Indels**<br>**(homozygous + heterozygous)** | 202 | 172 | 216 | 195 | 218 | 185 | 1,188 |
| **Hawaiian Indels**<br>**(homozygous)** | 6<br>(2.97%) | 6<br>(3.49%) | 6<br>(2.78%) | 20<br>(10.25%) | 7<br>(2.75%) | 6<br>(2.70%) | 49<br>(4.12%) |
| **Hawaiian SVs** | 59 | 49 | 68 | 58 | 95 | 58 | 387 |

We further analyzed each genome to understand the genomic distribution of SNPs and indels between each pair of Bristol and Hawaiian populations. We first analyzed the distribution of SNPs and indels per chromosome. For both genomes, the number of variant sites per chromosome roughly correlates with chromosome size, with the exception of the *X* chromosome (Figure 2). Between Bristol lineages, we identified 3,805 - 6,143 SNPs and 617 - 1,070 indels per chromosome, of which there are 33 - 60 SNPs and 25 – 40 indels that are homozygous sites on each chromosome (Figure 2A-B). In our comparison of Hawaiian genomes, we identified 934 - 2,185 SNPs and 1,889 - 2,523 indels per chromosome, of which there are 1 - 81 SNPs and 2 – 15 indels that are homozygous sites on each chromosome (Figure 2C-D). We then analyzed the spatial distribution of SNPs and indels across the length of each chromosome in 200 kb bins. Studies aligning the Bristol genome to the Hawaiian genome have demonstrated that much of the genomic variants exist on the highly-recombining, relatively gene-poor terminal “arm”-like domains that span several megabases at the end of each chromosome (C. elegans Sequencing Consortium 1998; Rockman and Kruglyak 2009; Thompson et al. 2015; D. Lee et al. 2021; Bush, Zachary D. et al. 2025; Kim et al. 2019). In our comparison of variant distributions in the two Bristol genomes, we found that every chromosome displays multiple intervals on one or both of the “arm”-like domains with a greater than average density of SNPs and/or indels (Figure 2E), although this pattern is not as striking as when aligning Bristol to Hawaiian (Thompson et al. 2015; Bush, Zachary D. et al. 2025). In contrast, for the two Hawaiian genomes, this broader pattern appeared much weaker across chromosomes, with the *X* chromosome not displaying any apparent biases for accumulation of variation in the terminal domains (Figure 2F). When we tested whether SNPs and indels were significantly enriched on the “arm” domains of each chromosome, Bristol variants were consistently enriched on the “arm” domains, while this pattern was less consistent on each chromosome for Hawaiian SNPs and indels (Supplemental Figure S2). Taken together, our data indicate that many variants are accumulating within inbred lab strains and a high fraction of heterozygous variant sites may exist between laboratory wild type *C. elegans* lineages.

### Genomic structural variations between independently cultivated lab strains

The constant development of new technologies that aid genome assembly, whole-genome alignments, and structural variant detection foster a new era of comparative genomics not limited in focus to the analysis of SNPs and indels. The rate of accumulation for SNPs and indels is known in *C. elegans* (Saxena et al. 2019), however, the rate of any type of SV per generation in *C. elegans* is largely unknown. Further, there is growing evidence for the impact of genomic SVs on genome evolution between the Bristol and Hawaiian isolates (Vergara et al. 2009; Maydan et al. 2010; Kim et al. 2019), yet very little is known about how many SVs have accumulated between different lineages of the Bristol or Hawaiian isolates in labs. To quantify and characterize SVs across each Bristol and Hawaiian genome, we utilized reciprocal whole-genome alignments between each pair of our *de novo* assemblies. For the DLW Bristol and VC2010 Bristol genomes, we found a total of 441 SVs between the two lineages (Table 2, Figure 3A). For the DLW Hawaiian and Kim Hawaiian genomes, we found a total of 387 SVs between the two lineages (Table 2, Figure 3A). Each chromosome in our Bristol comparison displayed between 71 and 78 SVs, and each chromosome in our Hawaiian comparison displayed between 49 and 95 SVs. Notably, the Hawaiian *X* chromosome (unlike the Bristol *X* chromosome) displayed a markedly lower fraction of the total SVs across the genome.

**Figure 3.**
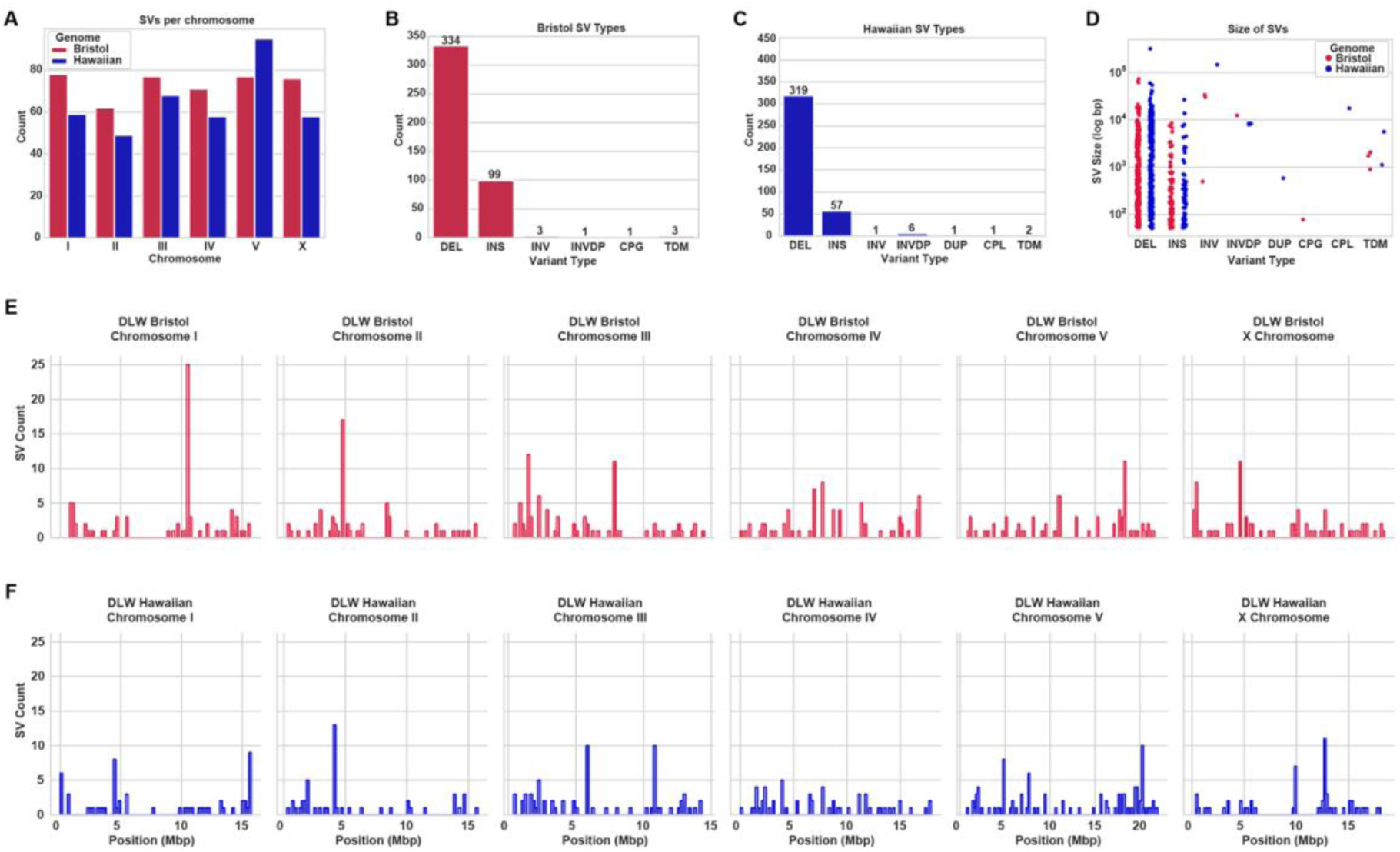
The genomic structural variants accumulated in laboratory lineages of C. elegans. **(A)** Bar chart showing the number of SVs on each chromosome in either the Bristol lineages or Hawaiian lineages of laboratory *C. elegans*. **(B)** Bar chart quantifying the number of each type of genomic SV between the DLW Bristol and VC2010 Bristol genome. **(C)** Bar chart quantifying the number of each type of genomic SV between the DLW Hawaiian and Kim Hawaiian genome. DEL = Deletion, INS = insertion, CPG = copy gain, CPL = copy loss, INV = inversion, INVDP = inverted duplication, and TDM = tandem repeat variation. **(D)** Strip plots showing the log-scale size distribution of all SVs called in either the Bristol or Hawaiian pairs of genomes. **(E)** Histograms showing the genomic distribution of SVs found on each chromosome between the DLW Bristol and VC2010 Bristol genomes. **(F)** Histograms showing the genomic distribution of SVs found on each chromosome between the DLW Hawaiian and Kim Hawaiian genomes. For E and F, SV counts are shown in non-overlapping 200 kb bins across the length of each chromosome.

Next, we wanted to identify the types of SVs that are present in each genome. Analysis of the alignments producing our SV calls revealed that deletions and insertions are the most common varieties of SVs present in each background. Laboratory lineages of the Bristol isolate have accumulated 334 deletion and 99 insertion SVs, with only eight SVs representing large copy-number variants, three inversions, and one inverted duplication (Figure 3B). Similarly, the Hawaiian genomes exhibit 319 deletion SVs and 57 insertion SVs in addition to11 SVs that are one copy loss variant, six inverted duplications, one inversion, one duplication, and two rearrangements in tandem-repeat regions (Figure 3C). In total, SVs account for 570 kb and 1,976 kb of variation between lineages of the Bristol and Hawaiian backgrounds, respectively. The size distribution of deletion and insertion SVs ranged from 50 bp up to hundreds of thousands of bases, with only four rearrangement SVs (not simple insertions or deletions) observed under 1 kb. The largest SV identified was a 317kb deletion on chromosome *IV* of the Kim Hawaiian lineage (Figure 3D). Lastly, the pattern of SV accumulation on the chromosome “arms” appears weaker in comparison to SNPs and indels, with many 200 kb intervals exhibiting zero SVs, and very few intervals displayed more SVs than the global average across all bins (Figure 3E-F). When we tested whether SVs were enriched on the chromosome “arms” versus the central domains, only chromosomes *II, III*, and the *X* chromosome in our Bristol lineage comparison and chromosomes *II* in our Hawaiian lineage comparison displayed this pattern (Supplemental Figure S2 and S3). Overall, these data identify the accumulation of hundreds of SVs over the course of individual lab cultivation of *C. elegans* wild type strains.

### Enrichment of SNPs, indels, and SVs in non-coding regions of the Bristol and Hawaiian genomes

Increasing evidence in multiple organisms has found that different regions of the genome accumulate variants at different rates, for which some can be attributed to local chromatin structure and the genomic function of a sequence (Makova and Hardison 2015; Ellegren, Smith, and Webster 2003; Zhang, Wang, and Andersen 2022; Schuster-Böckler and Lehner 2012; Wolfe, Sharp, and Li 1989; Hodgkinson and Eyre-Walker 2011). To identify features of the *C. elegans* genome that are particularly enriched (or depleted) of SNPs, indels, and SVs, we measured the overlap of variants with remapped sequence annotations. For the Bristol lineages, our analysis found that SNPs and indels in the Bristol lineages are moderately enriched in the introns of genes on the autosome (Log2(fold) values: SNPs 0.45-0.84; indels 0.50-1.02, all p-values < 0.05 by hypergeometric test; Figure 4A). Notably, we found a striking enrichment of indels in transcription factor binding sites on chromosome *V* (Log2(fold) value3.22, p-value < 0.05). In contrast to SNPs and indels, SVs are instead significantly enriched in the intergenic regions of all chromosomes except chromosome *I* (Log2(fold) values: 0.60-1.18, p-values < 0.05). For SNPs, indels, and SVs, we find a strong and significant depletion of variants in the coding sequences of genes on each chromosome (Log2(fold) values: SNPs −0.26 to −1.94, indels −1.16 to −3.84, and SVs −0.95 to −2.04; all p-values < 0.05). For the Hawaiian lineages, their genomes followed similar patterns to that of the Bristol lineages. Short sequence variants are moderately enriched in the introns of genes on the autosomes (Log2(fold) values: SNPs 0.67-1.04, indels 0.31-1.24; all p-values < 0.05; Figure 4B). SVs in the Hawaiian background are also enriched in intergenic sequences on all chromosomes (Log2(fold) values: 0.46-1.00, all p-values < 0.05). In contrast to our Bristol comparison, we did not find any consistent enrichment of SNPs and indels in regulatory sequences like enhancers, promoters, or transcription-factor binding sites, but SVs were enriched in enhancer sequences on chromosome *V* (Log2(fold) value: 2.08, p-value < 0.05). Instead, we found a much stronger depletion variants in coding sequences (Log2(fold) values: SNPs −1.54 to −4.23, indels −4.52 to −6.88, and SVs −0.94 to −2.08; all p-values < 0.05) than observed in Bristol lineages, which may reflect the much lower number of total SVs in the Hawaiian lineages. We did, however, find one striking example of accumulated variation in Hawaiian background. When we parsed our enrichment dataset by homozygous versus heterozygous variants, we found a strong enrichment of Hawaiian homozygous SNPs in coding sequences on chromosome *IV* (Log2(fold) value of 1.43, p < 0.05; Supplemental Figure S4. Finally, when we separately analyze the overlap of the variants of each genome binned into the “arms” versus central domains of chromosomes, we find that the magnitude of enrichment of SNP, indels, and SVs in non-coding sequences to be slightly higher in the central domains (Supplemental Figure S5). Taken together, our data in both Bristol and Hawaiian lineages find that nearly all variants reside in non-coding regions, and are highly depleted from exons and gene-regulatory sequences.

**Figure 4.**
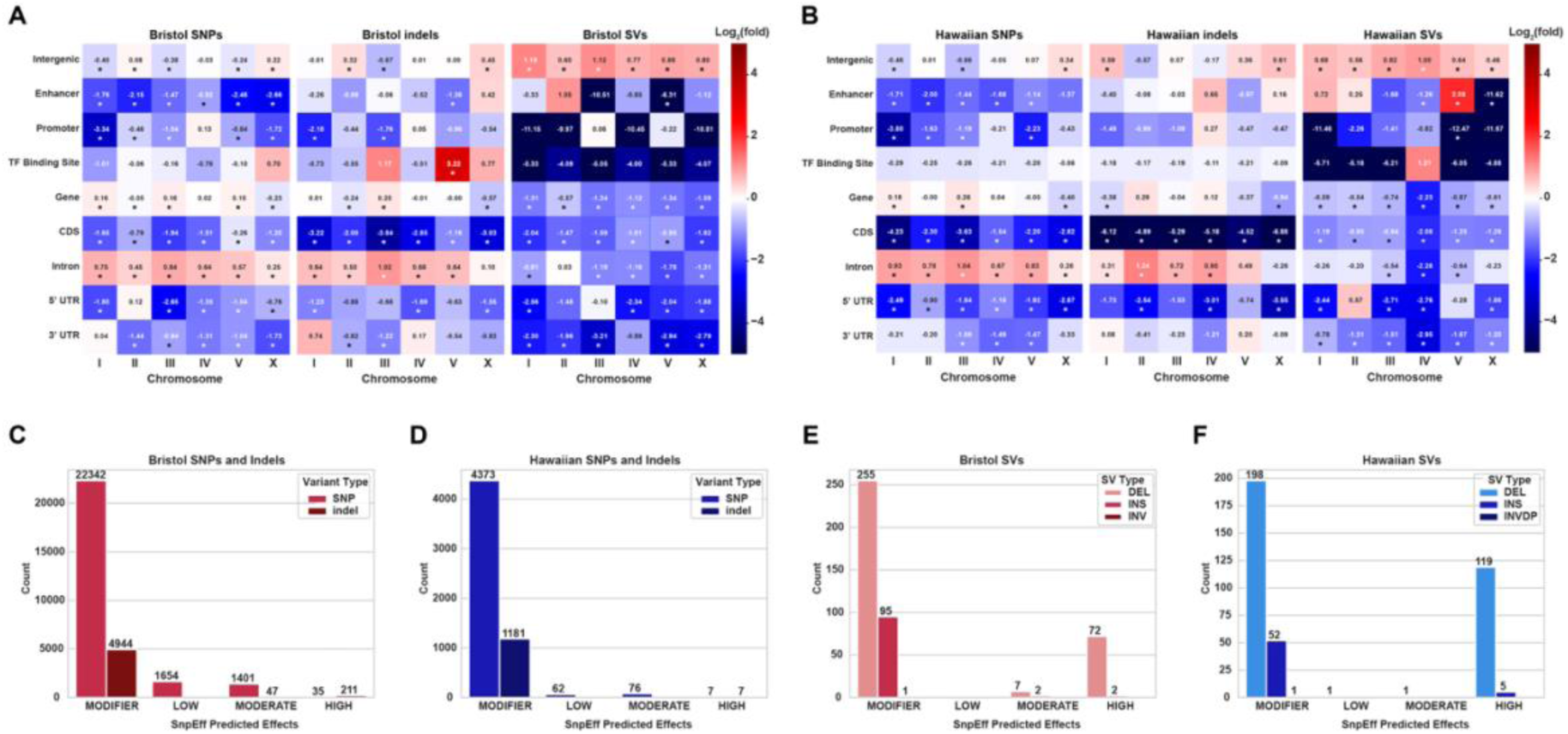
Genomic variants in laboratory *C. elegans* are enriched in non-coding regions. **(A)** Heatmap showing the log2(fold) enrichment or depletion of Bristol SNPs, indels, and SVs on each chromosome in genomic annotations. **(B)** Heatmap showing the log2(fold) enrichment or depletion of Hawaiian SNPs, indels, and SVs on each chromosome in genomic annotations. For A-B, asterisks under individual values indicate p-values calculated by the hypergeometric test are < 0.05. **(C)** Bar chart showing the predicted effects of Bristol SNPs and indels on gene function. **(D)** Bar chart showing the predicted effects of Hawaiian SNPs and indels on gene function. **(E)** Bar chart showing the predicted effects of Bristol SVs on gene function. **(F)** Bar chart showing the predicted effects of Hawaiian SVs on gene function.

To estimate the relative impacts of SNPs, indels, and SVs on the function of genes in our Bristol and Hawaiian comparisons, we performed analyses of the predicted functional changes to genes using SnpEff (Cingolani et al. 2012). From our analyses, we found that most predicted variant effects are low-consequence “modifier” effects, which are defined as: non-coding variants, variants affecting non-coding genes, or variants with unpredictable impact due to lack of empirical evidence. To contrast, “low”, “moderate”, and “high impact” variants include examples of synonymous mutations, missense/in-frame mutations, and frameshift/truncation mutations, respectively. In the Bristol genome, we found that SNPs and indels account for 246 high-impact effects and 1,448 moderate impacts to gene function (Figure 4C). Though SVs are much fewer in comparison to SNPs and indels, a greater fraction of SVs (17%) have moderate or high impact on gene function (Figure 4E). Similar to Bristol sequence variants, much of the SNPs and indels in our Hawaiian comparisons have few predicted impacts on gene function: 14 are high-impact SNPs/indels and 76 have moderate impacts (Figure 4E). SVs, on the other hand, frequently displayed high impacts on gene function in the Hawaiian background: 124/377 (∼33%) are predicted high impact, with large deletions representing 119 of these. In summary, while we found that the SNPs, indels, and SVs accumulate in non-coding regions of the genome at rates lower than expected by random chance, there is still potential for many of these variants (particularly SVs) to impact genome function and contribute to genome evolution.

## Discussion

Our examination of the inter-lab genetic drift among wild-type strains suggests that laboratory domestication of multiple *C. elegans* isolates has led to the accumulation of genomic variation between labs. Many of the variants we identified lie within non-coding regions of the genome such as intergenic sequences and regulatory sequences (*e.g.* transcription factor binding sites). Further, we identified large structural variations that could impact genomic-based analyses and other routine experimental practices such as primer design. Overall, our work reveals the genomic impact of long-term laboratory cultivation of wildtype *C. elegans* strains in different laboratories.

### Genomic divergence of laboratory wild type lineages

Earlier studies uncovering phenotypic and genetic variations between lab wild-type *C. elegans* strains indicated that there are likely many underlying large-scale genomic differences(Wicks et al. 2001; Sterken et al. 2015; Duveau and Félix 2012; Weber et al. 2010; McGrath et al. 2009; Rogers et al. 2003). Here we identify numerous SNPs, indels, and SVs between different lab lineages of the *C. elegans* wildtype isolates N2 Bristol and CB4856 Hawaiian. The total amount of genomic variation is at higher levels than predicted by earlier mutation accumulation studies (Denver et al. 2009; Saxena et al. 2019) The majority of total genomic variation is due to SVs, which have recently become a detailed subject of study (Thompson et al. 2015; Kim et al. 2019; Lee et al. 2021). Our genome assemblies of the DLW N2 Bristol and DLW CB4856 Hawaiian strains corroborate prior results indicating that genomic variation is enriched in the distal arm-like regions of chromosomes between these isolates (Thompson et al. 2015; D. Lee et al. 2021). Prior evolutionary genomic analyses suggest that both recombination in the arm-like regions of each chromosome and balancing selection likely shaped this landscape of sequence divergence across the 30,000-50,000 generations in which the N2 Bristol and CB4856 Hawaiian strains have been geographically isolated (Thomas et al. 2015; Kern and Hahn 2018). In contrast to comparisons between Bristol and Hawaiian genomes, we find that the distribution of variant sites across the “arm-like” regions versus “center-like” domains between lab lineages is not consistent across each chromosome. This result could indicate that in relatively short timescales (∼5,000-7,000 generations), selection for the accumulation of mutations in the arm-like regions, particularly in noncoding regions, is not sufficient to consistently eliminate sequence divergence away from the gene-dense chromosome centers. Further, we found that SNPs, indels, and SVs were highly enriched in introns and intergenic regions when aligning the genomes of laboratory strains. Selection against alleles that disrupt gene function may explain our observed reduced variation within coding regions, yet it remains possible that variants within some intergenic regulatory sequences (*e.g.* transcription factor binding sites, promoters, and/or enhancers) could produce emergent phenotypes. Thus, the accumulation of disruptive genomic changes within regulatory regions in the gene-dense centers of chromosomes may underpin some of the phenotypic differences observed in laboratory wild-type strains, such as variance in lifespan, fecundity, and social behavior (Gems and Riddle 2000; Andersen et al. 2014; Rogers et al. 2003).

### Potential impacts of accumulating structural variations

We detected many SVs with some up to hundreds of kilobases in size. Although these were often in intergenic regions of the genome, SVs of this size can be particularly disruptive to gene expression by impairing long range interactions between regulatory sequences such as promoters and enhancers that require specific three-dimensional genomic structures (Stranger et al. 2007; Hurles, Dermitzakis, and Tyler-Smith 2008). Further, the expression of eukaryotic genes relies on the splicing of introns out of pre-mRNAs, which is mediated by the spliceosome at specific recognition sites (Y. Lee and Rio 2015). Large insertions, deletions, and rearrangements within intronic sequences of genes can potentially disrupt pre-mRNA splicing and intronic copy number variations have been associated with variable gene expression in populations (Rigau et al. 2019). SVs within the introns of genes have also been shown to lead to the emergence of duplicated genes and give rise to functionally distinct paralogs (Xu et al. 2012). Therefore, the accumulation and prevalence of non-coding SVs cannot be ignored as they may lead to significant impacts on gene function and evolution between laboratory model systems.

Many experimental practices and genomic analyses rely on and utilize reference genomes. Aside from phenotypic consequences, the accumulation of undetected indels and SVs could be inhibitory to basic molecular biology techniques such as PCR or CRISPR if the target sequence is missing, disrupted, or rearranged. Further, SVs that are multi-kilobase gains, losses, and rearrangements of sequence, which may include entire genes, would be inhibitory to DNA sequencing workflows where the alignment of sequencing reads is necessary for downstream analyses. Our data identifying hundreds of kilobases or megabases of sequence divergence due to SNPs, indels, and SVs in laboratory populations of wild type laboratory strains suggests that usage of a *C. elegans* lineage with a recently published genome, or frequent return to cryogenically preserved *C. elegans* wild type stocks, may benefit molecular genetics and genomics experiments and analyses.

### Are Hawaiian *C. elegans* lineages exceptional in their variation?

We find that the total base pairs affected by genomic variation in CB4856 Hawaiian lineages is higher than our observations when aligning N2 Bristol lineages, which can be attributed to the presence of larger SVs. Although both strains passaged in labs have underappreciated amounts of variation, largely due to deletion/insertion SVs, the frequency of use for each strain should be considered. By far, the N2 Bristol strain is largely the standard wild type strain for laboratory research, and has been in use since the late 1960s and 1970s (Brenner 1974). The CB4856 Hawaiian isolate, however, has likely not been maintained or passaged as a live stock to the same extent in *C. elegans* labs worldwide as the N2 Bristol isolate. This difference in passaging frequency between the isolates could lead to lower frequencies of sequence variants (SNPs and indels) between individual laboratory lineages of CB4856 Hawaiian relative to individual laboratory Bristol strains. Notably, this lower frequency of sequence variants is despite the fact CB4856 Hawaiian *C. elegans* are a more “social” species with higher male mating frequencies in the population in contrast to N2 Bristol (Wegewitz, Schulenburg, and Streit 2008). This increased mating frequency could lead to increased heterozygosity and higher rate of sequence divergence between Hawaiian lineages in comparison to Bristol lineages. Further, the *Caenorhabditis* Genetics Center, an international repository and distributor of strains, returned to its 1995 working stock of CB4856 Hawaiian due to many labs reporting phenotypic abnormalities into 2013 and 2014 (CGC: https://cgc.umn.edu/strain/CB4856). Depending on how long each lab has passaged their lineage of the Bristol and Hawaiian strains, our account of SNPs and indels could be explained by findings in previous mutation accumulation studies (Denver et al. 2009). The rate of base substitutions (∼2.35 × 10^−9^ per site per generation) and indels (∼0.64 × 10^−9^ per site per generation) in the germline do not account for SV accumulation (Denver et al. 2009; Saxena et al. 2019), however, studies of human genomes support the lower counts of SVs observed in our *C. elegans* genomes (Nesta, Tafur, and Beck 2021). Why then, does there appear to be so much more genomic structural variation between lab lineages of the Hawaiian isolate? One possible source is that the Hawaiian genome experienced expansion of tandemly repeated regions (which include rDNA and regions of tandemly repeated TE sequences), which are known to mutate at much higher frequencies in yeast and humans (Fan and Chu 2007). Analysis of tandem repeat expansion between human individuals have shown that some tandemly repeated regions can vary in size from approximately 159.8 - 441.8 kb (Gondo et al. 1998). These remaining questions could be addressed, for example, with further sequencing of all laboratory *C. elegans* in question using the HiFi sequencing chemistry of PacBio combined with ultra-long Nanopore reads (Ichikawa et al. 2024).

## Conclusion

The generation of multiple independent *de novo* genome assemblies for both N2 Bristol and CB4856 Hawaiian isolates provides an excellent system to study genetic drift between laboratory model organisms. Additionally, identification and functional characterization of polymorphic sites and structural variations present between lab lineages of N2 Bristol and CB4856 Hawaiian may provide new insights into how pronounced phenotypic differences in the lifespan, feeding behavior, and reproductive fitness arise in modern lab-derived strains (Gems and Riddle 2000; Zhao et al. 2018). Future studies utilizing identical sequencing technologies and genome assembly methods for their comparisons will further illuminate the extent of genomic diversity between labs and allow for functional characterization of large genomic rearrangements.

## Data Availability Statement

The PacBio long-read and the Illumina short-read data used in this study to construct each DLW genome were previously submitted to the NCBI BioProject database (https://www.ncbi.nlm.nih.gov/bioproject/) under accession number PRJNA907379. PacBio long-read and Illumina short-read sequencing data for the VC2010 Bristol genome was accessed from the European Nucleotide Archive under the accession number PRJEB28388 (Yoshimura et al. 2019). PacBio long-read and Illumina short-read sequencing data for the Kim CB4856 Hawaiian genome was accessed from the NCBI Sequencing Read Archive under the BioProject accession number PRJNA523481 (Kim et al. 2019). Strains are available upon request.

## Acknowledgements

We thank N. Kurhanewicz, J. Conery, and other members of the Libuda lab for thoughtful critique and review of this manuscript during its preparation. We extend gratitude towards the University of Oregon Genomics and Cell Characterization Core Facility for the original library preps and sequencing of our DLW N2 Bristol and DLW CB4856 Hawaiian strains. We thank the CGC for providing multiple strains for this study (funded by National Institutes of Health P40 OD010440).

## Author Contributions

Z.D.B. devised strategies for genome assembly, assembled the contigs, filled gaps, and performed all variant calling and related analyses. A.F.S.N, D.D., and K.J.H. helped conceive this study and develop strategies for genome assembly. D.E.L. helped conceive this study, led discussions for the comparison of different genomes, procured funding, and coordinated work. All authors contributed to the manuscript, with most of the writing by Z.D.B. and D.E.L.

