## Supplemental Figures for "Genomic sequence and structural variations accumulating between laboratory lineages of wild type *C. elegans*": supplemental_figure_S1.pdf

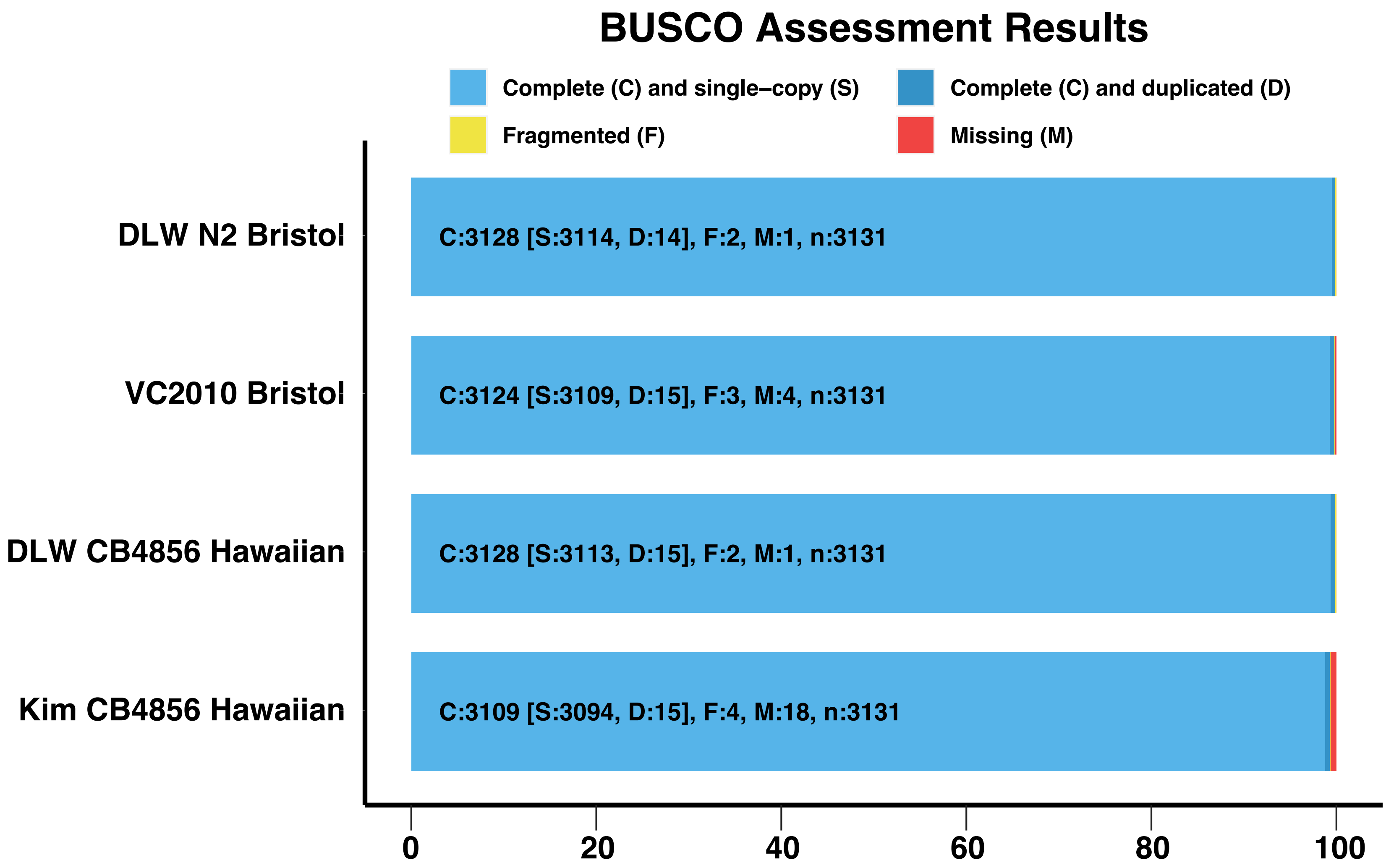


Supplemental Figure S1. BUSCO analysis of Bristol and Hawaiian genome assemblies. The presence of orthologous genes from metazoan and nematode lineages are shown foreach genome assembly. Each orthologous gene analyzed is depicted as either Complete (C, blues), Fragmented (F, yellow), or Missing (M, red). Complete orthologs are then further categorized as single-copy (S, light blue) or duplicated (D, dark blue). The "nematoda odb10" database was downloaded and used as the reference set of orthologous genes.
