## Supplemental Figures for "Genomic sequence and structural variations accumulating between laboratory lineages of wild type *C. elegans*": supplemental_figure_S2.pdf

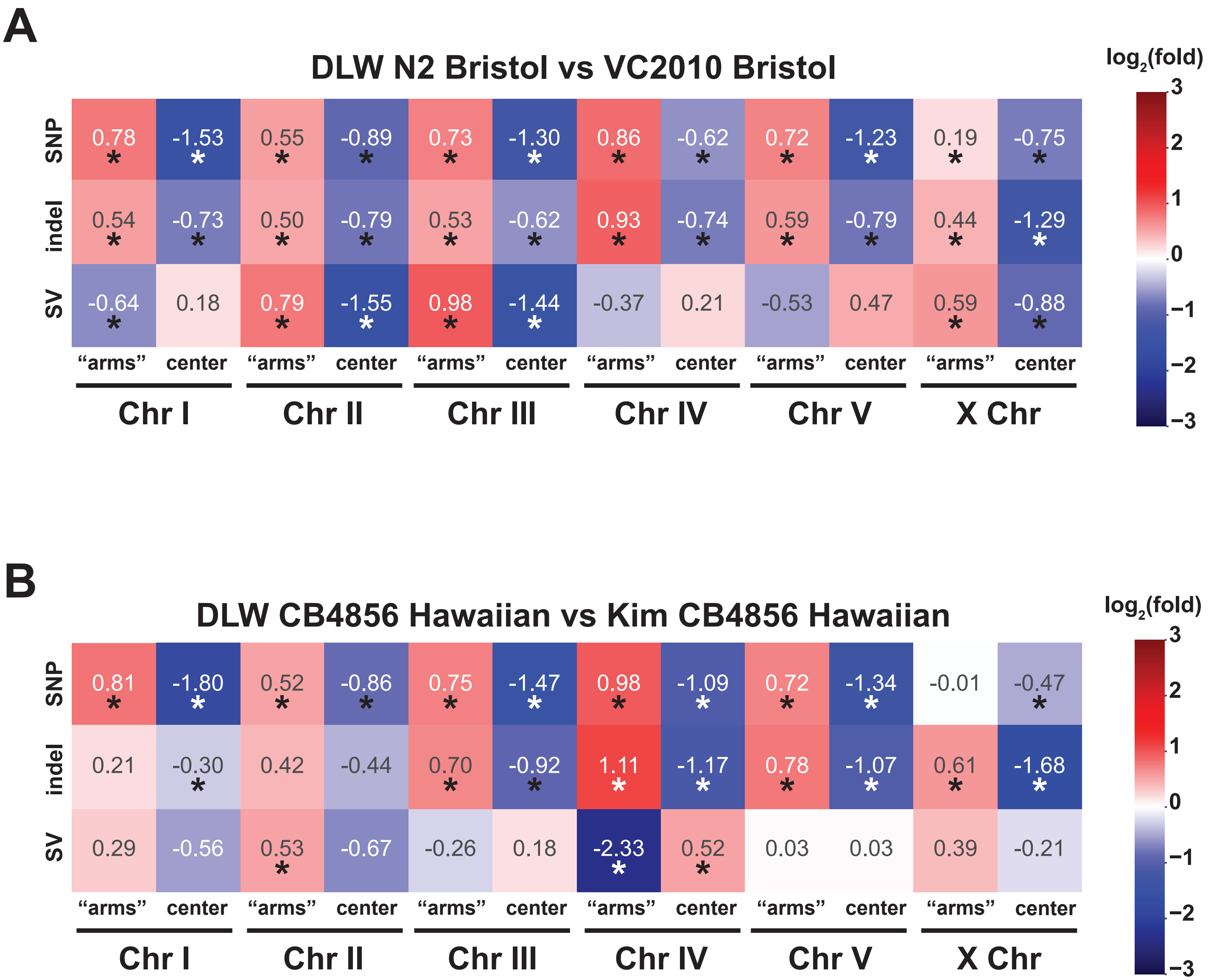


Supplemental Figure S2. The enrichment of variants within the "arms" versus central domain of each chromosome. (A) Heatmap showing the Log2(fold) enrichment or depletion of each variant type in the "arms" (combined) or central region of each chromosome in our comparison of the Bristol genomes. (B) Heatmap showing the Log2(fold) enrichment or depletion of each variant type in the "arms" (combined) or central region of each chromosome in our comparison of the Hawaiian genomes. Asterisks below values indicate that the degree of overlap between variants and each annotation is significantly different ($p < 0.05$) that the distribution of overlaps generated by 10,000 simulated null distributions.
