## Supplemental Figures for "Genomic sequence and structural variations accumulating between laboratory lineages of wild type *C. elegans*": supplemental_figure_S3.pdf

**A**

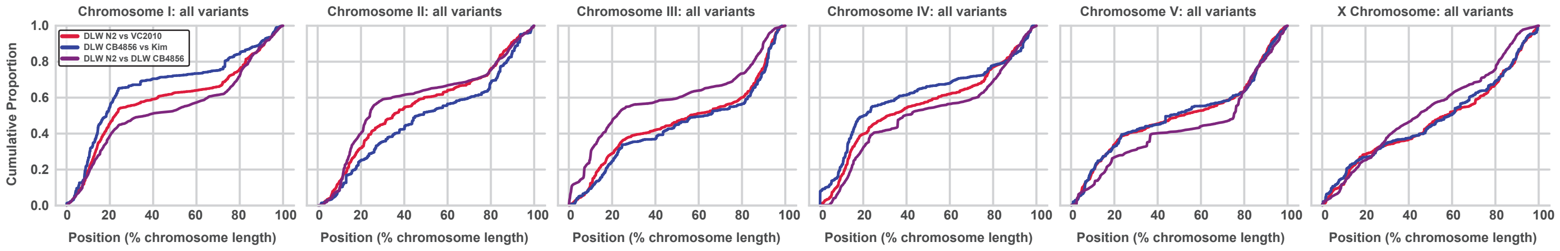


| Variant Distribution Comparisons | Chromosome | | | | | |
|---|---|---|---|---|---|---|
| | *I* | *II* | *III* | *IV* | *V* | *X* |
| **Lab Bristols vs Lab Hawaiians** | *** | *** | ** | *** | n.s. | n.s. |
| **Lab Bristols vs "Bristol : Hawaiian"** | *** | *** | *** | *** | *** | *** |
| **Lab Hawaiians vs "Bristol : Hawaiian"** | *** | *** | *** | *** | *** | *** |

**B**

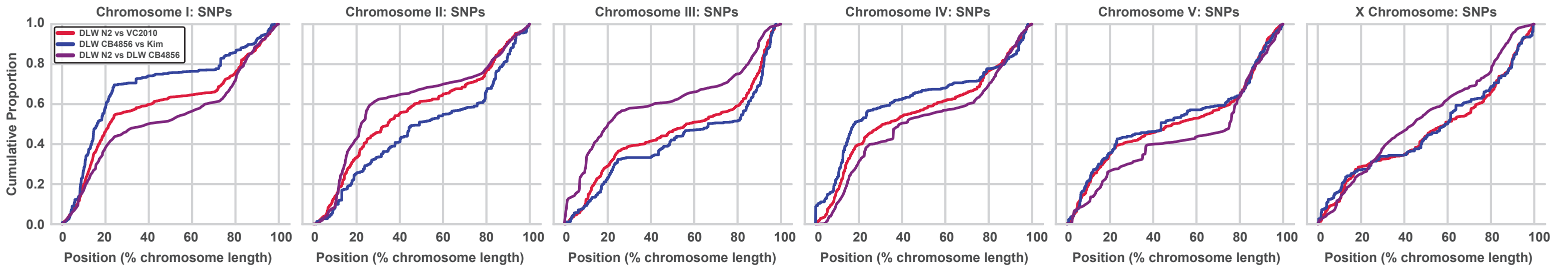


| SNP Distribution Comparisons | Chromosome | | | | | |
|---|---|---|---|---|---|---|
| | *I* | *II* | *III* | *IV* | *V* | *X* |
| **Lab Bristols vs Lab Hawaiians** | *** | *** | *** | *** | n.s | n.s |
| **Lab Bristols vs "Bristol : Hawaiian"** | *** | *** | *** | *** | *** | *** |
| **Lab Hawaiians vs "Bristol : Hawaiian"** | *** | *** | *** | *** | *** | *** |

**C**

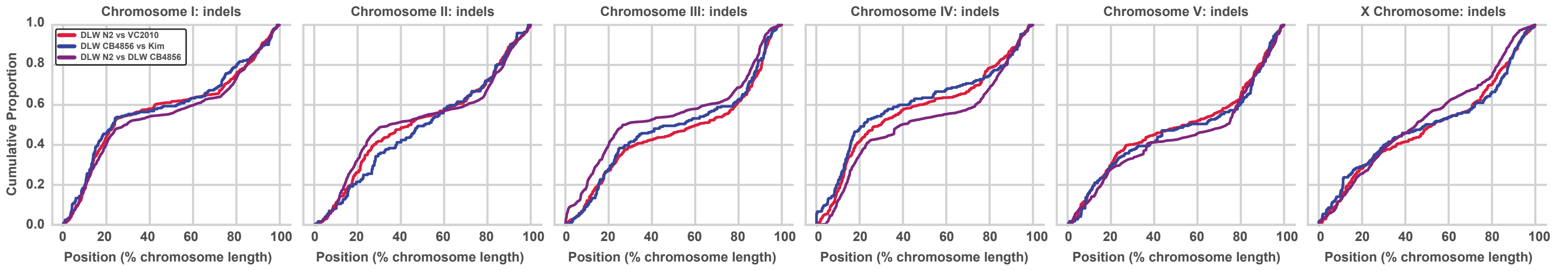


| Indel Distribution Comparisons | Chromosome | | | | | |
|---|---|---|---|---|---|---|
| | *I* | *II* | *III* | *IV* | *V* | *X* |
| **Lab Bristols vs Lab Hawaiians** | n.s. | * | n.s | n.s | n.s | n.s |
| **Lab Bristols vs "Bristol : Hawaiian"** | ** | *** | *** | *** | *** | *** |
| **Lab Hawaiians vs "Bristol : Hawaiian"** | n.s. | *** | *** | *** | n.s. | ** |

**D**

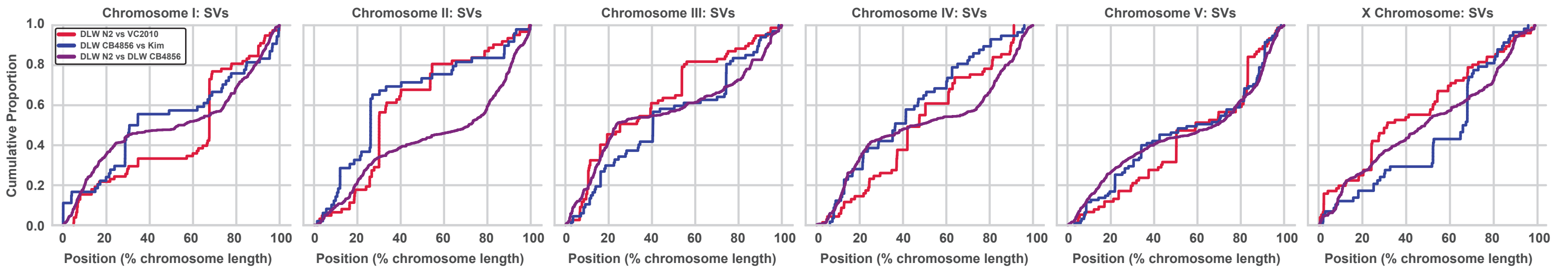


| SV Distribution Comparisons | Chromosome | | | | | |
|---|---|---|---|---|---|---|
| | *I* | *II* | *III* | *IV* | *V* | *X* |
| **Lab Bristols vs Lab Hawaiians** | * | *** | n.s | n.s | n.s | ** |
| **Lab Bristols vs "Bristol : Hawaiian"** | ** | *** | ** | ** | * | n.s |
| **Lab Hawaiians vs "Bristol : Hawaiian"** | n.s. | *** | * | *** | n.s. | ** |

Supplemental Figure S3. Comparisons of variant distributions in recently diverged laboratory lineages versus diverged wild isolates. (A) ECDF plots showing the distributions of all variants on each chromosome. (B) ECDF plots showing the distributions of SNPs on each chromosome. (C) ECDF plots showing the distributions of indels on each chromosome. (D) ECDF plots showing the distributions of SVs on each chromosome. For each panel, the distributions of variants comparing the DLW Bristol vs VC2010 genomes are shown in red, the distributions of variants comparing DLW Hawaiian to the Kim Hawaiian genome are shown in blue, and the variant distributions observed when comparing the N2 Bristol genome to the CB4856 Hawaiian genome are shown in purple. Below each set of ECDF plots is a table indicating the significance of p-values determined by two-sample KS tests for goodness of fit when comparing the distribution of variants in laboratory lineages versus the distribution of variants in wild isolates.
