## Supplemental Figures for "Genomic sequence and structural variations accumulating between laboratory lineages of wild type *C. elegans*": supplemental_figure_S4.pdf

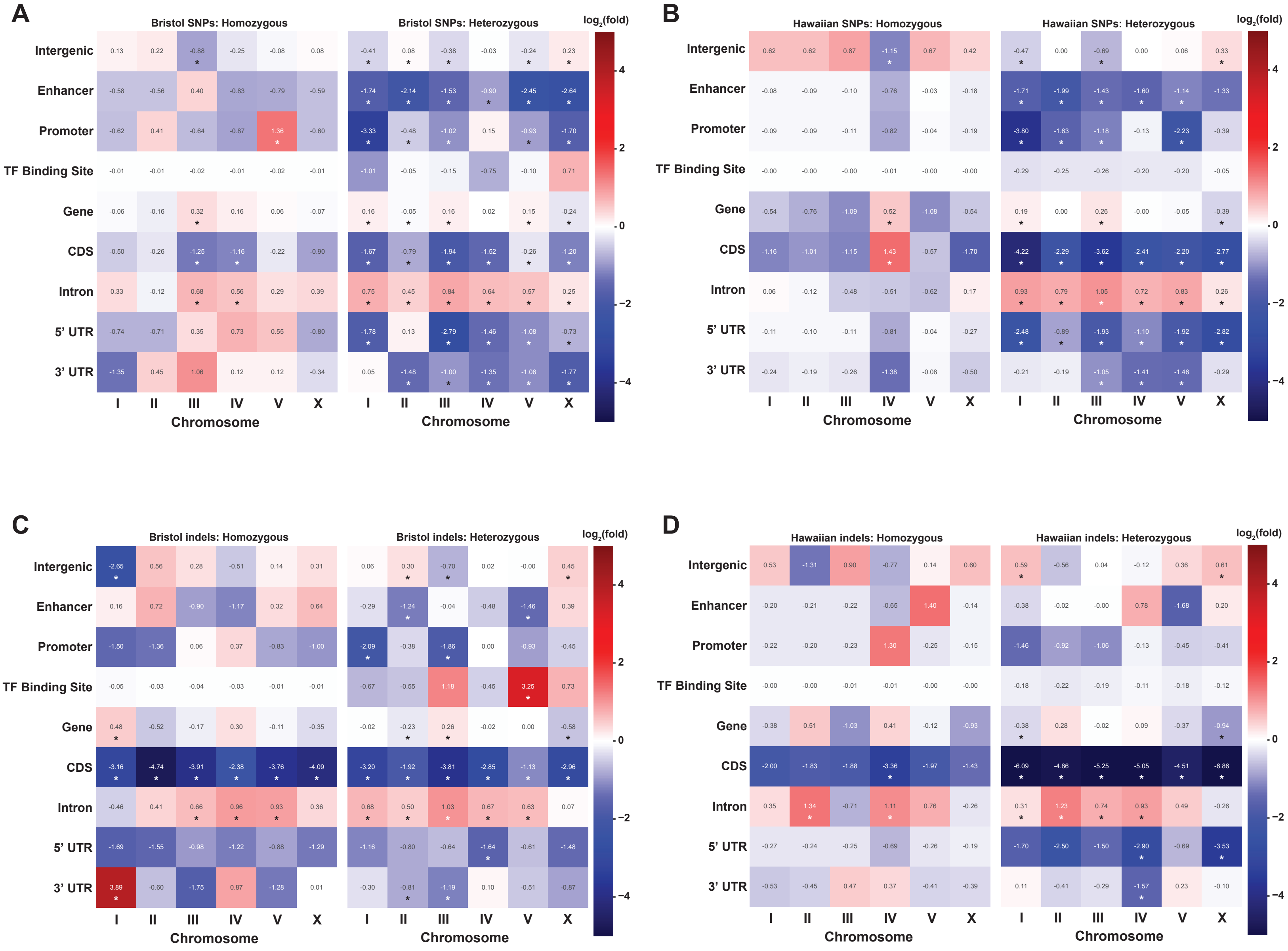


Supplemental Figure S4. The enrichment of variants in sequence annotations parsed by heterozygous versus homozygous sites across each genome. (A) Heatmap showing the Log2(fold) enrichment or depletion of homozygous and heterozygous Bristol SNPs in gene annotations on each chromosome. (B) Heatmap showing the Log2(fold) enrichment or depletion of homozygous and heterozygous Hawaiian SNPs in gene annotations on each chromosome. (C) Heatmap showing the Log2(fold) enrichment or depletion of homozygous and heterozygous Bristol indels in gene annotations on each chromosome. (D) Heatmap showing the Log2(fold) enrichment or depletion of homozygous and heterozygous Hawaiian indels in gene annotations on each chromosome. Asterisks below values indicate that the degree of overlap between variants and each annotation is signicantly different ($p < 0.05$) that the distribution of overlaps generated by 10,000 simulated null distributions.
