## Supplemental Figures for "Genomic sequence and structural variations accumulating between laboratory lineages of wild type *C. elegans*": supplemental_figure_S5.pdf

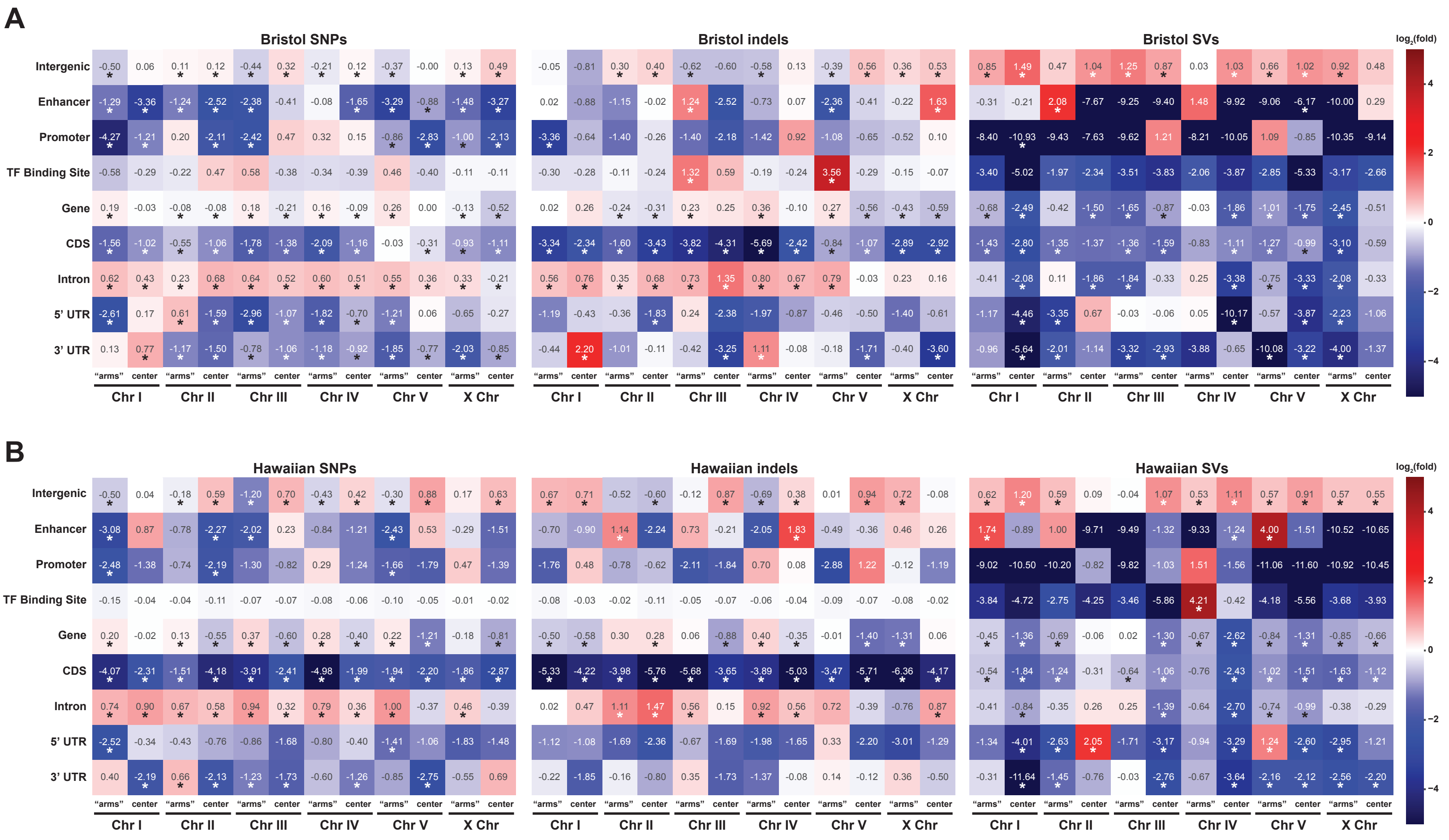

Supplemental Figure S5. The enrichment of variants in sequence annotations on the arms versus central domains of each chromosome. (A) Heatmap showing the Log2(fold) enrichment or depletion of Bristol SNPs, indels, and SVs in gene annotations on each chromosome. (B) Heatmap showing the Log2(fold) enrichment or depletion of Hawaiian SNPs, indels, and SVs in gene annotations on each chromosome. Asterisks below values indicate that the degree of overlap between variants and each annotation is signicantly different ($p < 0.05$) that the distribution of overlaps generated by 10,000 simulated null distributions.
